# Ectopic, hepatic GLP-1R agonism enhances the weight loss efficacy of GLP-1 analogues

**DOI:** 10.1101/2025.04.01.646483

**Authors:** Jonathan D. Douros, Megan Capozzi, Aaron Novikoff, Jacek Mokrosinski, Barent DuBois, Joseph Stock, Rebecca Rohlfs, Mikayla Anderson, Dominika J. Jedrzejcyk, Svend Poulsen, Erik Oude Blenke, Tomas Dago, Kasper Huus, Peder L. Nørby, Sune Kobberup, Marita Rivir, Joyce Sorrell, Stephanie Mowery, Daniel J. Drucker, David A. D’Alessio, Jonathan E. Campbell, Timo D. Müller, Diego Perez-Tilve, Patrick J. Knerr, Brian Finan

## Abstract

**Objective:** Unimolecular triagonists drive substantial weight loss in patients with obesity (PwO) by engaging the glucagon-like peptide 1 (GLP-1) and glucose dependent insulinotropic polypeptide (GIP) receptors to reduce food intake (FI) and the hepatic glucagon (Gcg) receptor to enhance energy expenditure (EE). However, their development has been challenged by deleterious cardiovascular (CV) effects including increased heart rate (HR), elongated QTc, and arrhythmia mediated by GcgR agonism. GLP-1R monoagonists on the other hand improve both obesity and CV outcomes with negligible effects on EE. We sought to imbue peptide GLP-1R agonists with an EE enhancing effect by combining them with ectopic GLP-1R expression and agonism in hepatocytes.

**Methods:** We used an attenuated adenovirus (AAV) to induce the expression of a functional, liver-specific GLP-1R combined with traditional peptide agonist treatment to drive greater body weight loss via reduced energy intake and increased energy expenditure.

**Results:** Agonism of the ectopic GLP-1R with either semaglutide, a low internalization GLP-1R agonist (Sema584), or a dual GLP-1R/GIPR agonist in wild-type (WT) diet induced obese (DIO) mice led to enhanced EE and improved weight loss compared to agonist treatment alone.

**Conclusions:** This represents a novel mechanism for achieving polypharmacy to treat obesity.

**Highlights:**

- A Glp1r encoding AAV induces expression of a functional receptor mouse livers.
- Endogenous GLP-1R does not mediate semaglutide clearance.
- Ectopic GLP-1R mediates semaglutide clearance.
- Ectopic, hepatic Glp1r plus semaglutide enhances weight loss in mice.
- Ectopic, hepatic Glp1r plus a dual incretin agonist enhances weight loss in mice.

## Introduction

Triple agonists of the GLP-1R, GIPR, and GcgR drive profound weight loss in PwO [1-4]. While the full complement of mechanisms underlying these effects remains to be fully characterized, it is clear that pharmacologic activation of central GLP-1R[5] and GIPR[6; 7] populations reduces FI, while hepatic GcgR agonism promotes EE [8; 9] in rodents. Unlike their mono- and dual-incretin receptor agonist predecessors, which improve CV function [10-13], development of GcgR activating compounds has been challenged by the increased HR and arrhythmia induced by chronic GcgR agonism [2; 3; 14]. Therefore, we sought to enhance the safe and efficacious weight lowering capacity of GLP-1R- and dual GLP-1R/GIPR agonists by augmenting their FI lowering effect with an ability to enhance energy expenditure. To accomplish this, we took advantage of intrinsic similarities and differences in GLP-1R and GcgR. Both receptors couple to Gαs and utilize cAMP as a second messenger, but only GcgR is robustly expressed in hepatocytes where its pharmacologic activation drives energy expenditure. However, the chronotropic effect of GcgR agonism is linked, at least in rats, to action in the sinoatrial node [15]. Thus, we hypothesized that ectopically expressing and subsequently activating a GLP-1R in hepatocytes would lead to enhanced EE and improved weight loss with no additional increase in HR.

## Methods

### Attenuated adenovirus (AAV)

The AAV serotype 8 construct was generated by Vector Builder with the *mGlp1r* construct downstream of a liver specific promotor

### LNP

Lipid nanoparticle formulations were prepared using a NanoAssemblr Ignite microfluidic mixer (Precision Nanosystems) by mixing at a 3:1 volumetric ratio (aqueous:ethanol) and 12 mL/min total flow rate. Lipid solutions were made in pure ethanol at a molar ratio of [50:10:38.5:1.5] for [ionizable lipid:DSPC:cholesterol:DMG-PEG2000] at a total lipid concentration of 12.5 mM. The ionizable lipid (heptadecan-9-yl 8-((2-hydroxyethyl)(8-(nonyloxy)-8-oxooctyl)amino)octanoate) was synthesized as previously described [16]. DSPC, cholesterol and DMG-PEG2000 were purchased from Sigma-Aldrich. mRNA solution was prepared using nuclease free reagents with final buffer concentration of 25 mM acetate (Sigma-Aldrich) and corresponding to a N/P ratio of 6 when mixed with the lipid excipients.

After microfluidic mixing, LNPs were transferred to 10 kDa MWCO Slide-A-Lyzer Dialysis Cassettes (ThermoFisher Scientific) and dialyzed overnight against 20 mM Tris (Sigma-Aldrich), 8 % (w/vol) sucrose (Sigma-Aldrich), pH 7.4. After dialysis, the formulation was sterilized via filtration through a 0.2 µm syringe filter and concentrated by centrifuging at 1000 x g in Amicon Ultra centrifugal filter unit with a 100 kDa MWCO (Millipore Sigma). After concentration, the formulation was diluted to a final concentration of 0.3 mg/mL mRNA with 20 mM Tris, 8 % (w/vol) sucrose, pH 7.4. LNPs were characterized by dynamic light scattering using a Zetasizer Nano (Malvern Panalytical); the particle diameter (Z-Ave) was 70.7 nm with a PDI of 0.11. mRNA concentration and encapsulation efficiency measurement were performed using Quant-iT RiboGreen reagent (Invitrogen); the encapsulation efficiency was 95.3 % in the final formulation.

### mRNA design

*Mus musculus* glucagon-like peptide 1 receptor (*mGlp1r*) (NM_021332.2) coding sequence was codon optimized for expression in murine cells. mRNA was designed to start with GGGA prior to 5’UTR derived from a TOP gene. Two stop codons were added downstream of the *mGlp1r* coding sequence to ensure translation termination. A 70 nt-long poly(A) tail was added to the end of albumin gene 3’UTR.

### mRNA synthesis

mRNA was enzymatically, *in vitro* transcribed from double-stranded, PCR-amplified DNA template. Single-stranded DNA template including T7 promoter was chemically synthesized (IDT) and PCR-amplified with SuperFi II PCR Master Mix (ThermoFisher Scientific) with reverse primer containing 70 nt-long dT repeated fragment (IDT). Amplified IVT template was transcribed using a T7 enzyme mix containing T7 polymerase, RNase Inhibitor and inorganic pyrophosphatase (ThermoFisher Scientific). Uridine (U) was substituted with modified uridine analog, N1-methyl-pseudouridine (N1meψ) (TriLink). The reaction was incubated 4 h at 37 °C with 350 rpm mixing. Following IVT, capping mixture containing Vaccinia Capping Enzyme, Vaccinia mRNA Cap2’-O-Methyltransferase, S-adenosyl-methionine (SAM), and GTP (ThermoFisher Scientific) was added, and reaction was incubated 1 h at 37 °C with 350 rpm mixing. IVT template was removed from reaction by digestion with RNase-free DNase I (ThermoFisher Scientific). mRNA was purified applying Carboxylic Acid Dynabeads for RNA Purification (ThermoFisher Scientific) according to manufacturer’s protocol. mRNA quality, integrity and purity were assessed by capillary electrophoresis and A260/280 ratio respectively.

### Peptides

All peptide agonists utilized in these studies including native GLP-1, semaglutide, Sema584, and the dual GLP-1R/GIPR agonist were produced according to protocols that have been described in detail elsewhere[8]. Briefly, peptides were constructed using automated Fmoc-base solid-phase peptide synthesis. Following completion of the peptide backbone, acylation with a fatty acid-based protractor was achieved using an orthogonal protecting group strategy on the appropriate lysine sidechain. The peptide was then cleaved from resin, purified by reversed-phase high-performance liquid chromatography, and lyophilized to dryness. Dry peptide was dissolved in a buffer containing 50 mM phosphate, 70 mM sodium chloride, and 0.05% Tween 80 for use in animal studies.

### In vitro assays

The *in vitro* assays used to characterize GLP-1R internalization induced semaglutide, Sema584, and the dual GLP-1R/GIPR agonist have been described in detail elsewhere [17]. Briefly, the GLP-1R internalization bioluminescent resonance energy transfer assay was established by measuring loss of baseline resonance energy transfer between an intracellular plasma membrane marker GFP-CAAX and hGLP-1R-RLUC8 or hGIPR-RLUC8 following ligand administration. Transfections were performed using 500 ng of GFP-CAAX DNA and 300 ng of the respective RLUC8-tagged GPCR DNA per well in a 6-well plate.

To assess the ability of the *Glp1r* LNP to induce GLP-1R expression *in vitro*, 50 ng LNP was added to 5000 BHK CRE-Luc reporter cells in 24 microtiter plate well replicates. Following 18 hours incubation at 37°C and 5% CO2, a 12-point titration of semaglutide was added to the cells in duplicates, resulting in a final volume per well of 25 µL. After 4 hours incubation at 37°C and 5% CO2, 12.5 µL detection reagent (Steady-GLO, E2510, Promega) was added to each well and luminescence was detected for each well using a multimode plate reader (EnVision 2105, PerkinElmer).

### Animal studies

All animal studies were performed at University of Cincinnati in accordance with approved IACUC protocols. Mice with DIO were given *ab libitum* access to water and a 58% fat, high-sugar diet (D12331, Research Diets) for at least 12 weeks animals were housed 3-4 per cage, exposed to a controlled 12 h/12 h light–dark cycle at room temperature (22 °C).

### GLP-1R KO mouse studies

For studies with DIO GLP-1R KO mice [18], male animals (mean BW 39.7g) were randomized to treatment groups (n = 3/group) based on starting body-weight. Animals were treated with *mGLP-1R* expressing AAV (10^11^ or 10^12^ GC/mouse) or control AAV (10^12^) at day −7 via tail vein injection. Animals were allowed to recover from day −7 to 0. On day 0 the semaglutide challenge was performed (described below). Body weight and food intake was assessed from day 0 to day 7 with no active treatment. From day 13 to 33 animals received daily SC injections of semaglutide (3nmol/kg) in all groups; body weight and food intake was assessed. On day 27 body composition was assessed (described below). Animals were sacrificed and tissues were collected on day 34 for qPCR assessment of GLP-1R mRNA expression (described below). Body weight loss on day 33 was correlated with hepatic GLP-1R mRNA expression.

### Proglucagon knockout mice

The proglucagon (ProG) KO mouse line has been validated in previous reports [19; 20]. Briefly, ProG KO and wild-type control mice were treated with either control or *mGlp1r* expressing AAV 7d prior to experiments. We then performed either IPGTT, oral glucose tolerance tests, or mixed nutrient tolerance tests as previously reported [21; 22].

### Semaglutide challenge

The semaglutide challenge in DIO GLP-1R mice was performed by fasting mice for 6h prior to the experiment, then administering semaglutide (3nmol/kg) via SC injection at t = 0. Blood glucose was measured from the tail vein via handheld glucometer over a 6h time course.

### Body composition

Whole body fat and lean mass were measured using nuclear magnetic resonance (EchoMRI, TX).

### Weight loss studies in wild type mouse studies

Male C57Bl6 mice were randomized and evenly distributed to test groups (n = 8 per group) according to body weight. Animals received tail vein injection of the control AAV or *mGLP-1R* expressing AAV at a dose of 10^12^ GC per mouse on day −7 for each study. Semaglutide, Sema584, and dual GLP-1R/GIPR agonist treatment began on day 0 for each study. Treatments were administered via daily SC injection for the duration and dosage indicated in Figures 2, 3, and 4. Dose escalation regimens for each peptide were determined based on previously published studies [8]. Body weight and food intake was measured every other day throughout the study.

**Figure 1:**
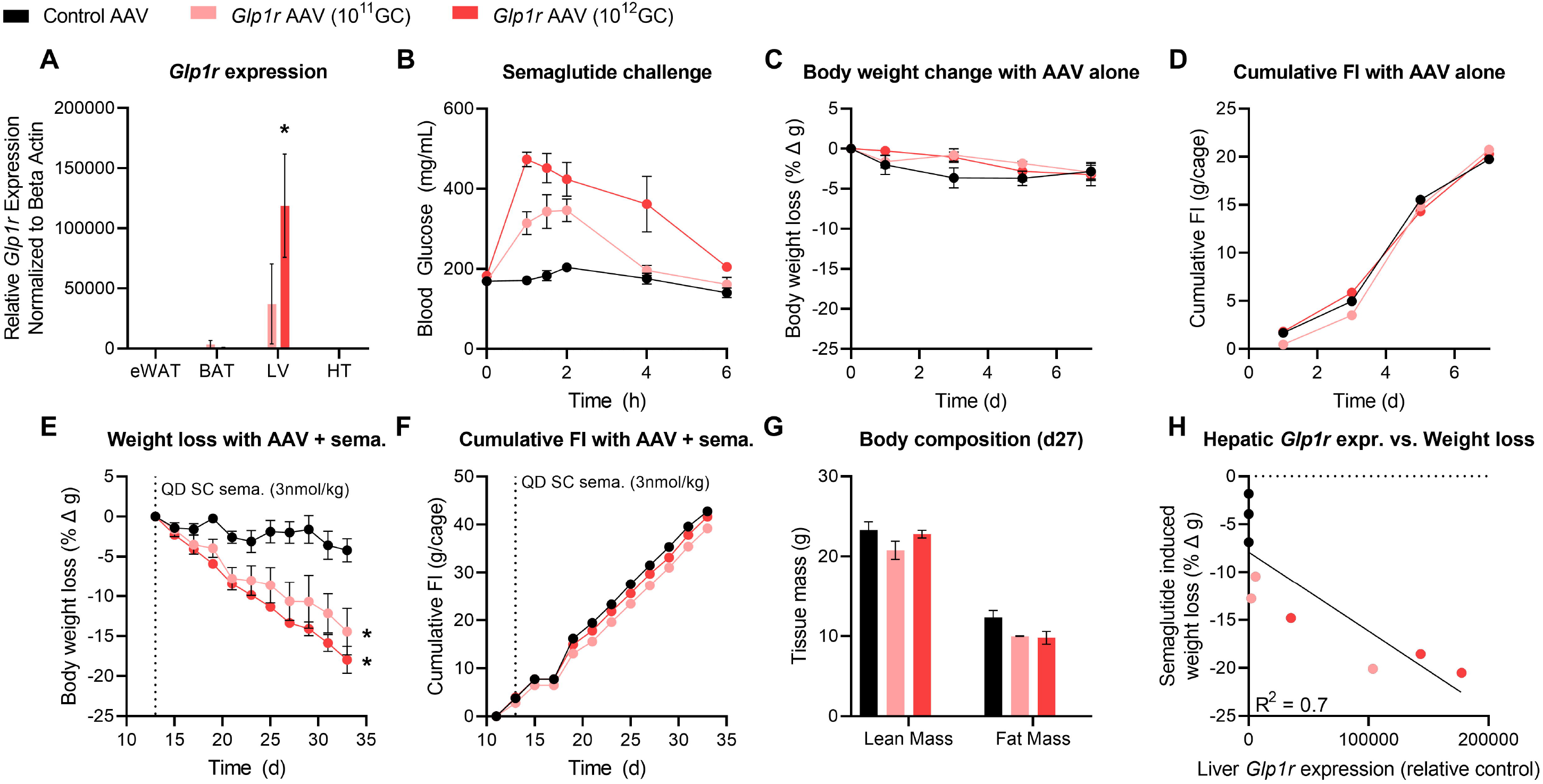
A *Glp1r* encoding AAV induces expression of a functional receptor in the liver of DIO GLP-1R knockout mice. Global GLP-1R knockout mice on HFD were dosed I.V. with a control antibody (black bars or circles) mouse *Glp1r* expressing AAV at 2 doses (10^11^ in light red, 10^12^ in red; genome copies: GC) per mouse. (A)*Glp1r* gene expression relative to beta-actin in the epididymal white adipose tissue (eWAT), brown adipose tis (BAT), liver (LV), and hypothalamus (HT) (B) Blood glucose after a single semaglutide dose (3nmol/kg) (C,D) Body weight change and cumulative food intake (FI) in response to the *Glp1r* expressing AAV alone dosed on d0. (E,F,G) Body weight change, cumulative FI, and body composition (d27) in response to the *Glp1r* expressing AAV + semaglutide (QD, SC 3 nmol/kg started d13) (H) Correlation between hepatic *Glp1r* expression and semaglutide mediated weight loss All data are presented as mean ± SEM. * indicates a p-value ≤ 0.05 compared to control AAV treated animals

**Figure 2:**
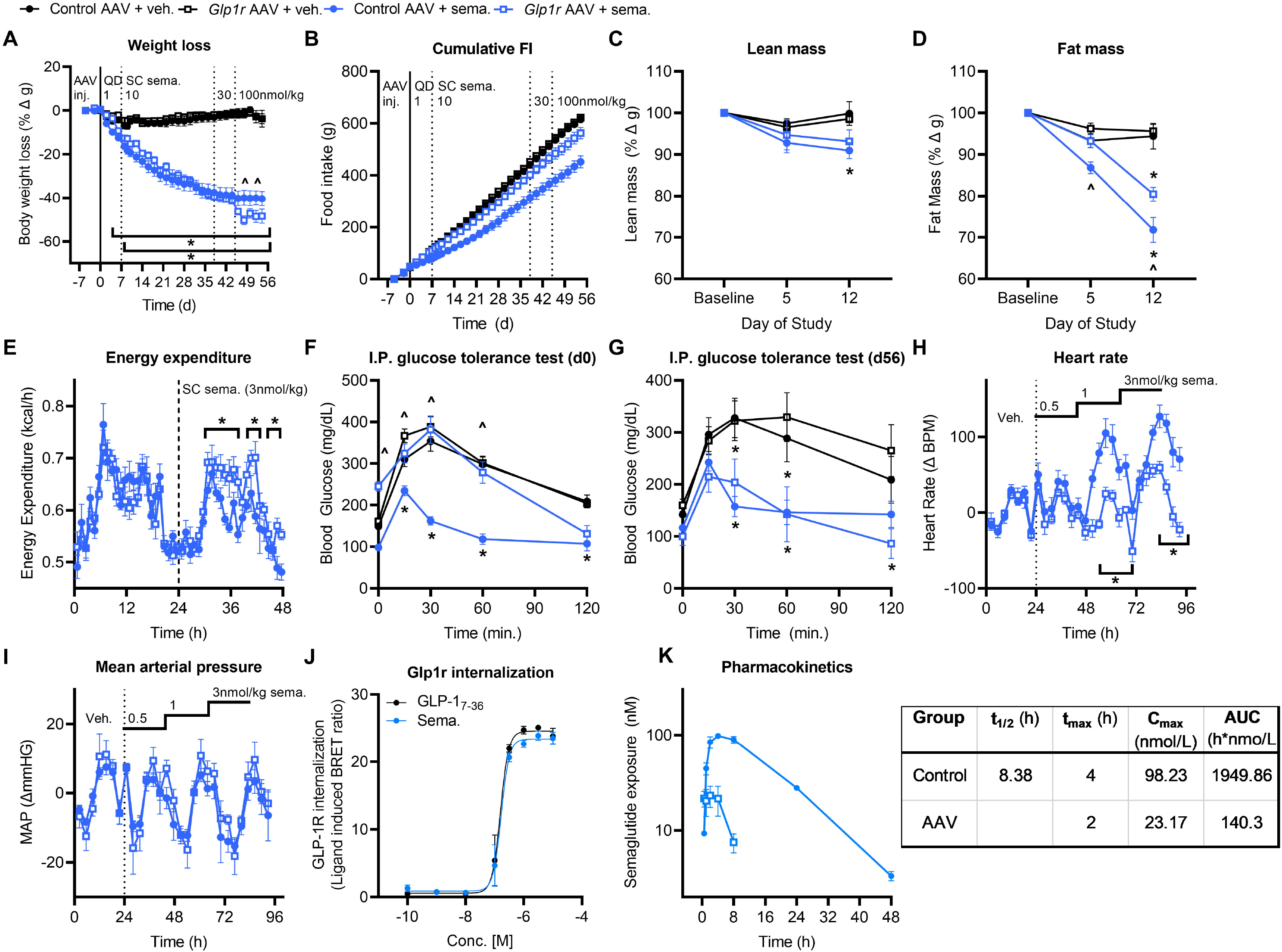
Coupling ectopic, hepatic *Glp1r* expression and semaglutide treatment enhances weight loss in DIO mice. WT DIO mice were dosed with a *Glp1r* expressing AAV (10^12^ GC) on −7d then with semaglutide (dose escalated from 1-100 nmol/kg) from d0 to 56. (A,B) BW loss and cumulative FI (C,D) Lean and fat mass d 0, 5 and 12 (E) EE during an acute semaglutide injection (3 nmol/kg) (F,G) Blood glucose during IPGTTs performed on d0 (F) and d56 (G). (H,I) Heart rate and mean arterial pressure assessed in control AAV and *Glp1r* AAV treated mice in response to semaglutide (0.5, 1, and 3nmol/kg SC doses) (J) *In vitro* GLP-1R internalization induced by native GLP-1 or semaglutide (K) *In vivo* PK profile for semaglutide in WT DIO mice dosed with either control (closed circles) or *Glp1r* expressing AAV (10^12^ GC, open circles) All data are presented as mean ± SEM. * indicates a p-value ≤ 0.05 compared to respective control animals ^^^ indicates a p-value ≤ 0.05 between control AAV + semaglutide and *Glp1r* AAV + semaglutide groups Notations under/over time points indicate significant difference at that time point determined by 2way ANOVA with Tukey multiple comparisons test. Notations to the right of the data indicate a significant group effect determined by 2way ANOVA with Tukey multiple comparisons.

**Figure 3:**
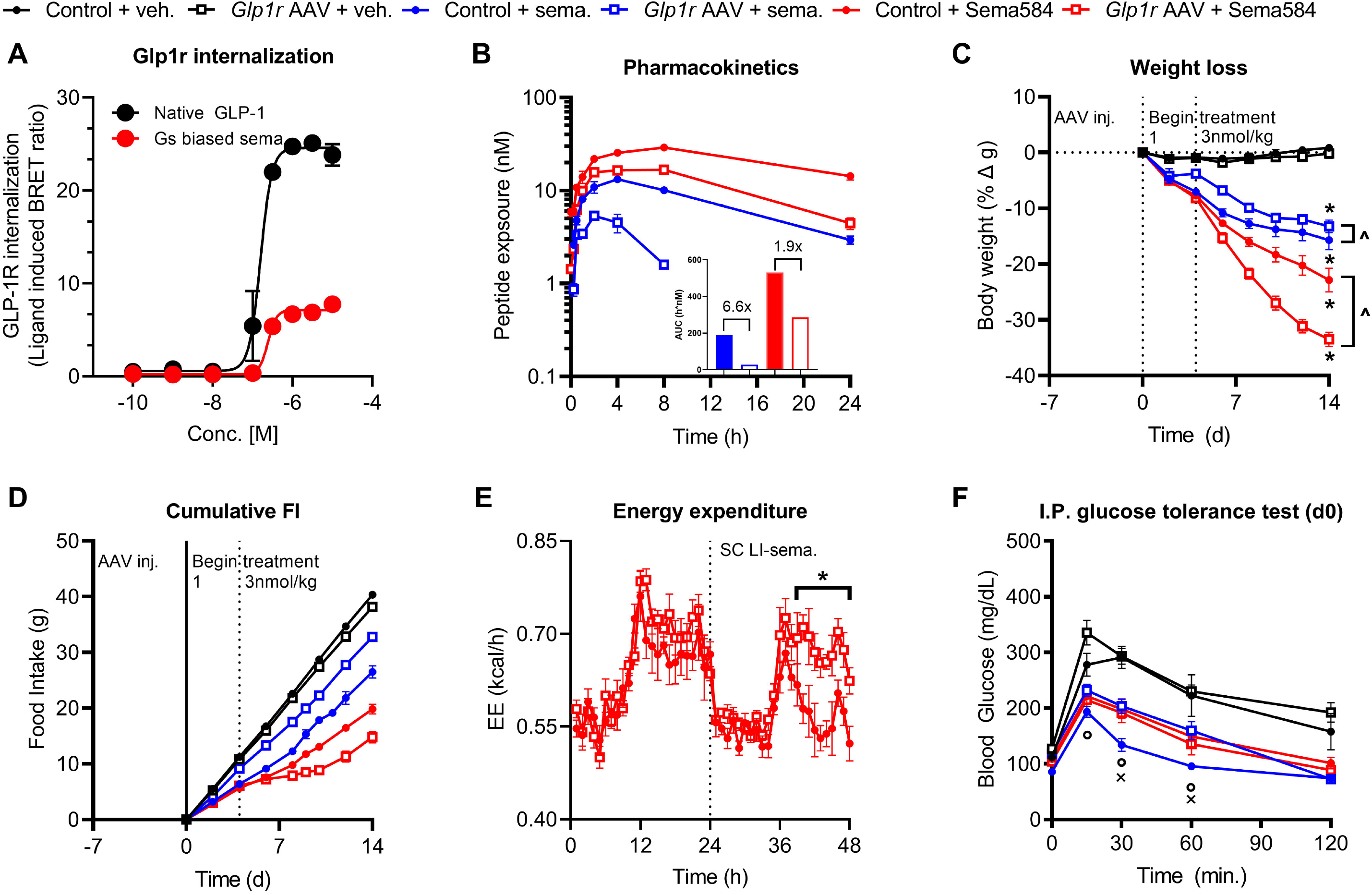
GLP-1R agonist that exhibits limited receptor internalization can obviate the PK liability of hepatic *Glp1r* expression. (A)*In vitro* GLP-1R internalization induced by native GLP-1 or LI-sema. (B)*In vivo* PK profile for semaglutide or LI-sema. in WT DIO mice dosed with either control or *Glp1r* expressing AAV (10^12^ GC) (C,D) Weight loss and cumulative FI during a 14d treatment regimen with either vehicle, semaglutide or a LI-sema. in WT DIO mice dosed with either control or *Glp1r* expressing AAV (10^12^ GC) (E)EE after acute treatment with either semaglutide or a LI-sema. in WT DIO mice dosed with either control or *Glp1r* expressing AAV (10^12^ GC) (F) Blood glucose during an I.P. glucose tolerance test performed on d0 All data are presented as mean ± SEM. *indicates a p-value ≤ 0.05 compared to respective control animals ^^^ indicates a p-value ≤ 0.05 between groups as indicated ^ο^ indicates a p-value ≤ 0.05 for all groups compared to their respective controls ^x^ indicates a p-value ≤ 0.05 between control AAV + semaglutide and *Glp1r* AAV + semaglutide groups Notations under/over time points indicate significant difference at that time point determined by 2way ANOVA with Tukey multiple comparisons test. Notations to the right of the data indicate a significant group effect determined by 2way ANOVA with Tukey multiple comparisons.

**Figure 4:**
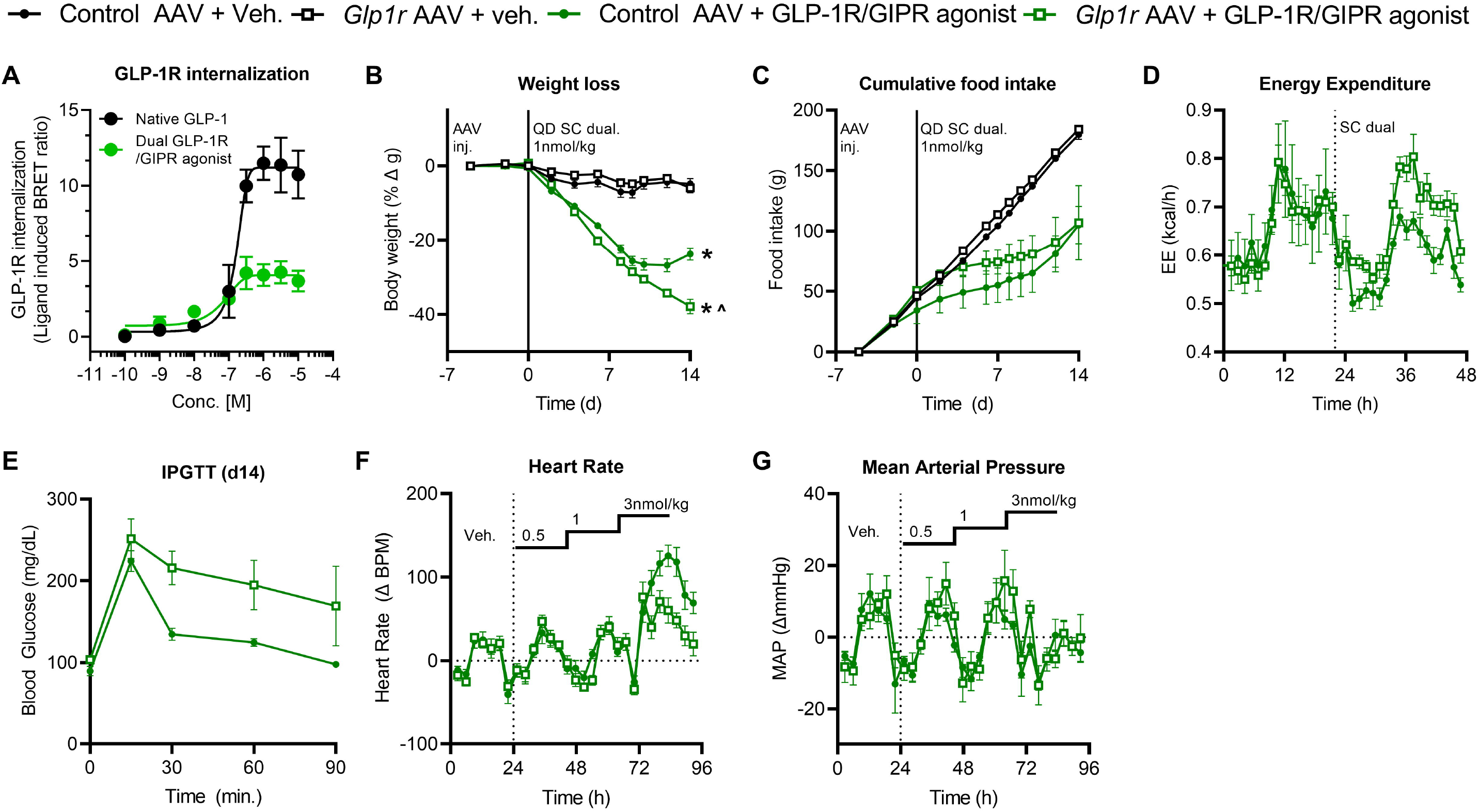
The weight lowering efficacy of a dual incretin agonist is enhanced in the presence of ectopic, hepatic *Glp1r* expression. (A) *In vitro* GLP-1R internalization for native GLP-1, semaglutide, LI-sema., and a dual GLP-1R/GIPR agonist (B,C) Weight loss and cumulative FI during a 14d treatment regimen with either vehicle or a dual GLP-1R/GIPR agonist at 1 nmol/kg in control or *Glp1r* AAV treated mice (D) Acute EE response to a dual GLP-1R/GIPR agonist in AAV and *Glp1r* AAV treated mice (E) Blood glucose during an IPGTT on d14 of the study (F,G) Acute HR and MAP response to a dual GLP-1R/GIPR agonist in AAV and *Glp1r* AAV treated mice All data are presented as mean ± SEM. *indicates a p-value ≤ 0.05 compared to respective control animals ^^^ indicates a p-value ≤ 0.05 between control AAV + GLP-1R/GIPR agonist and *Glp1r* AAV + GLP-1R/GIPR agonist groups Notations under/over time points indicate significant difference at that time point determined by 2way ANOVA with Tukey multiple comparisons test. Notations to the right of the data indicate a significant group effect determined by 2way ANOVA with Tukey multiple comparisons.

### IPGTT

Mice were fasted at the onset of the light phase for 6h prior to the experiment. Treatment compounds were injected at t = −1h to allow for SC absorption of the compound into circulation prior injection of a glucose bolus (2g/kg, ip, 20% dextrose in 0.9% NaCl). Blood glucose was measured via handheld glucometer (Abbot Freestyle) over a 2h time course as outlined.

### Indirect calorimetry

Hepatic GLP-1R expression was induced by AAV injection in DIO WT studies as outlined above. Energy expenditure was assessed as previously reported [23]. Briefly, animals were singly housed in the indirect calorimetry systems (*TSE Phenomaster/Labmaster, TSE Systems, Chesterfield, MO, USA)* for up to 14d and allowed to acclimate for up to 5d before measurements began, including QD SC vehicle injections and monitoring baseline body weight and food intake. During the study, mice were either monitored during SC administration of QD compounds, or given single injections and measured during the washout period. Body weight and food intake were also monitored.

### Cardiovascular radiotelemetry

Male C57Bl6 mice (8-10week old) were anesthetized with isoflurane and surgically implanted with pressure transmitters (*PA-C10, DSI*), with the catheter inserted into the left carotid artery and the body of the transmitter between the outer skin layer and abdomen wall in the left flank of the mouse. After recovering, they were singly housed and monitored in their home cages. Mice were given control and experimental AAV virus via tail vein injections at least 7 days prior to CV study. Transmitters were turned on via magnets and SC vehicle injections were administered 2 hours after the onset of the light phase for baseline measurements. At t=24h, mice were grouped if needed and SC compound injections began. Stressors were allowed only from 9:00 AM to 11:00 AM during the light phase, with continuous sampling throughout the day. Measurements of pulse pressure, arterial pressures, and heart rate values were extracted with the DataQuest A.R.T. 4.1 software and averaged into 3-hour bins for analysis.

### Gene expression

Gene expression was assessed in the liver, hypothalamus, white adipose tissue, and brown adipose tissue of GLP-1R KO mice given either control or *mGLP-1R* expressing AAV via qPCR protocols as reported elsewhere. The tissue samples were processed in a tissue lyser and RNA was extracted from the samples following the RNeasy MiniKit (*Qiagen, Cat. No. 74106*) and DNase I treatment (*Qiagen, Cat. No. 79256)*. Using the iScript kit (*Bio-Rad, 1708890*), cDNA was synthesized. The Taqman probes used are against Glp1r (*ThermoFisher, Mm00445292_m1*) as the gene of interest, and Beta Actin (*ThermoFisher, 4352341E)* as the endogenous control. Plates were read with the QuantStudio 5 system, and the resulting values were used to calculate the ΔΔCT, normalized to Beta Actin.

### Pharmacokinetics

For pharmacokinetic studies in wild-type and GLP-1R KO mice, animals were dosed subcutaneously with semaglutide or Sema584glutide as described above. Pharmacokinetic profiles were assessed according to previously reported methods [24]. Briefly, plasma concentration-time profiles were analysed by a non-compartmental method (Pharsight Phoenix WinNonLin v.6.4). The terminal half-life (t_1/2_), maximum plasma concentration (C_max_), time for maximum plasma concentration (T_max_), and AUC from zero to last (AUC_0-t_) were determined. Criteria for estimation of t_1/2_ were at least three concentration-time points in the terminal phase not including Cmax, with an R^2^ ≥□□0.85.

### LC/MS bioanalysis

Plasma concentrations of 0113-5840 and semaglutide were determined by liquid chromatography-tandem mass spectrometry (LC-MS/MS) using a multiple reaction monitoring method. Briefly, plasma proteins were precipitated by mixing plasma samples with 6 volumes of methanol containing internal standard in micronic 1.5mL microcentrifuge tubes, followed by centrifugation for twenty minutes at 13,000xg. The supernatant was transferred to a 96-well plate and diluted with water containing 0.1% formic acid, and mixed thoroughly. Diluted samples were injected into the LC-MS/MS system. The chromatographic separation was performed on a Waters I-class LC system using a Waters Acquity UPLC BEH C18 column (1.0 mm x 50 mm, 1.7 μm) with gradient elution of 0.1% formic acid in water (mobile phase A) and 0.1% formic acid in acetonitrile (mobile phase B) at a flow rate of 0.3 mL/min with a column temperature of 60□°C. The mass spectrometric detection was performed on a Thermo Scientific Quantiva triple quadrupole system with electrospray ionization in positive ion mode.

## Results

We utilized a *mGlp1r*-encoding AAV8 with a human α-antitrypsin promotor dosed via tail-vein injection to drive liver specific expression (*Glp1r* AAV). In lean WT mice, the *Glp1r* AAV treatment alone did not affect body weight (BW; Supplemental figure 1A,E) but did worsen glucose control (Supplemental figure 1B,C,D). This trend toward glucose dysregulation is mediated by endogenous proglucagon products activating the ectopic GLP-1R in hepatocytes, and recapitulates what is observed with acute glucagon administration (Supplemental figure 1F,G,H). While this outcome in lean mice is deleterious, the model does not represent the pathophysiology we aim to treat. Additionally, these data serve as an indication that the *Glp1r* AAV drives expression of a functional GLP-1R on the liver that mimics GcgR.

We next treated DIO GLP-1R KO mice with the *Glp1r* AAV at 2 doses (10^11^ or 10^12^ genome copies; GC) per mouse (Figure 1A-H). Gene expression for *Glp1r* was dose-dependently elevated by *Glp1r* AAV treatment at in the liver, but not in the white adipose tissue (eWAT), brown adipose tissue (BAT), and hypothalamus (HT; Figure 1A). Blood glucose was not acutely reduced in control AAV treated GLP-1R KO mice after a single injection of GLP-1R monoagonist semaglutide (3nmol/kg), as expected given the lack of GLP-1R in these animals (Figure 1B). However, semaglutide acutely increased blood glucose in GLP-1R KO mice given the *Glp1r* AAV, consistent with the ectopic, hepatic expression of a functional GLP-1R that mimics GcgR agonism (Figure 1B). The *Glp1r* AAV groups exhibited no change in BW or FI with AAV treatment alone (Figure 1C,D). On initiation of once-daily subcutaneous semaglutide injections (d13, Figure 1E,F) both *Glp1r* AAV treated groups lost BW, but showed little reduction in FI compared to control AAV treated animals, indicative of an increase in energy expenditure. Fat and lean mass declined consistently in the *Glp1r* AAV treated groups compared to controls (Figure 1G). The BW change induced by semaglutide treatment correlated well to hepatic *Glp1r* gene expression (Figure 1H). We also utilized a lipid nanoparticle (LNP) encapsulating an mRNA construct to successfully drive *in vitro* (Supplemental figure 2A) and *in vivo* (Supplemental figure 2B,C) GLP-1R expression as we feel this modality has more translational value. Semaglutide-mediated blood glucose excursion (Supplemental figure 2B,C) in was increased in DIO GLP-1R KO mice treated with this *Glp1r* expressing LNP construct, although at a reduced magnitude compared to the AAV construct. These data support the conclusion that a GLP-1R ectopically expressed in the liver can functionally mimic the effects of GcgR agonism with respect to glucose excursion and FI-independent weight reduction in rodents.

Expanding from these findings, we tested whether the pharmacologic effects of triagonists to reduce FI and enhance EE can be replicated by cotreatment with the *Glp1r* AAV and semaglutide in DIO WT mice. The animals were dosed with either control or *Glp1r* AAV (10^12^ GC) on day −7 and began treatment with either vehicle or semaglutide (dose escalation from 1 to 100 nmol/kg) on day 0. Unexpectedly, the semaglutide and semaglutide + *Glp1r* AAV groups show similar BW, fat mass, and lean mass loss (Figure 2A,C,D) at doses of 1 to 30 nmol/kg semaglutide (Figure 2A; d0-45). It is noteworthy that there is minimal FI reduction in the semaglutide + *Glp1r* AAV during this period compared to control AAV or *Glp1r* AAV alone (Figure 2B; d0-45). However, at 100 nmol/kg semaglutide, we observe a stark decline in BW concomitant with reduced FI for the semaglutide + *Glp1r* AAV group compared to the group treated with semaglutide alone. This is collectively suggestive that lower doses of semaglutide (1 to 30 nmol/kg) drive BW reduction through two different mechanisms in the *Glp1r* AAV and control AAV treated mice mice. Mice treated with control AAV + semaglutide alone achieve BW loss via reduced FI (i.e. GLP-1 acting as GLP-1) whereas mice treated with *Glp1r* AAV + semaglutide appear to experience BW loss via increased EE (i.e. GLP-1 mimicking glucagon). This is supported by the observation that semaglutide + *Glp1r* AAV treated mice exhibit increased EE compared to semaglutide treatment alone in an acute setting (Figure 2E). We observe semaglutide treatment alone acutely increases HR in mice with no effect on mean arterial pressure (MAP; Figure 2H,I), in keeping with previous reports. Intriguingly, semaglutide + *Glp1r* AAV reduces this effect. We hypothesized that the modified pharmacology of semaglutide at lower doses (i.e. increased EE, no effect on FI, IPGTT, or HR at doses ≤ 30 nmol/kg) is due to ectopic GLP-1R mediated drug clearance. *In vitro* GLP-1R internalization mediated by semaglutide is comparable to native GLP-1 (Figure 2F) suggesting a mechanism for ectopic GLP-1R to drive drug clearance. Endogenous GLP-1R populations do not facilitate a significant degree of semaglutide clearance (Supplemental figure 3), however the additional pool of GLP-1Rs in the liver created by *Glp1r* AAV treatment facilitate a significant increase in semaglutide clearance compared to control groups. From these data, we conclude that ectopic hepatic GLP-1R agonism via semaglutide in DIO mice can drive enhanced EE and additional BW loss compared to GLP-1R agonism alone without deleterious increases in HR. However, the full potential of this approach is obviated by ectopic GLP-1R mediated drug clearance.

Because there is an apparent clearance of semaglutide via internalization of the ectopic, hepatic GLP-1R we hypothesized that a GLP-1R agonist which displays reduced ligand-mediated receptor internalization would circumvent this mechanism. We selected a semaglutide-like peptide agonist that induces a low degree of receptor internalization (Sema584; Figure 3A) for testing in rodents. Sema584 showed a 1.9-fold reduction in circulating exposure in *Glp1r* AAV treated mice compared to controls, whereas semaglutide displayed a 6.6-fold reduction (Figure 3B). Repeated dosing of Sema584 in DIO mice pre-treated with *Glp1r* AAV resulted in superior BW and, critically, FI reduction (Figure 3C,D) compared to controls, while semaglutide treatment recapitulated our findings from previous studies (Figure 2). Accordingly, EE was also acutely elevated by Sema584 in *Glp1r* AAV treated animals compared to control mice in a separate experiment (Figure 3E). Blood glucose during an IPGTT on d0 was similar between control AAV and *Glp1r* AAV treated mice given Sema584 (Figure 3F). This data supports the hypothesis that a GLP-1R monoagonist with low ligand-mediated receptor internalization (e.g. Sema584) can limit the target mediated clearance mechanism presented by ectopic GLP-1R expression. Furthermore, combining GLP-1R agonists like Sema584 with ectopic GLP-1R expression provides greater BW reducing efficacy than the GLP-1R monoagonist alone by engaging multiple mechanism(s) of action (MoA) including FI reduction and EE enhancement.

Finally, we tested the hypothesis that ectopic GLP-1R agonism can be leveraged to improve the efficacy of dual GLP-1R/GIPR agonists as a surrogate for triagonism. We began by demonstrating that our previously published dual incretin receptor agonist [8] exhibits low GLP-1R internalization *in vitro* (Figure 4A). These data are in keeping with other dual GLP-1R/GIPR agonists including tirzepatide [25] and suggest our tool compound will be able to avoid the pharmacokinetic (PK) liabilities of the highly internalizing agonist semaglutide. The dual GLP-1R/GIPR agonist was able to induce greater BW reduction in DIO WT mice pre-treated *Glp1r* AAV compared to control AAV treated animals (Figure 4B). The dual GLP-1R/GIPR agonists induces is a transient reduction in FI between *Glp1r* AAV and control AAV treated groups (Figure 4C), which is a typical, yet poorly understood, effect of triagonist treatment in rodents [8; 9]. These data suggest a mechanism by which activation of liver GPCRs act to reduce FI, but also do not necessarily rule out the hypothesized action of glucagon to reduce FI via a central nervous system mechanism. Additionally, we see a significant elevation in EE during acute dual GLP-1R/GIPR agonist treatment in the *Glp1r* AAV treated animals compared to controls (Figure 4D). We note a deterioration of glucose control in *Glp1r* AAV treated mice after 14d of treatment compared to controls (Figure 4E). Finally, there is no additional increase in HR (Figure 4F) and no changes in MAP (Figure 4G) when combining *Glp1r* AAV with the dual receptor agonist at a dose (1nmol/kg) which drives superior BW loss. This indicates that the increased energy expenditure is not inherently linked to the CV liabilities of GcgR agonism as has been hypothesized [26].

## Discussion

These results lead to several conclusions that advance our goal of developing poly-pharmacologic approaches to treat obesity. The first is that functional GPCRs can be expressed ectopically to facilitate a poly-pharmacy platform. The *Glp1r* AAV and LNP/mRNA modalities employed in these studies drive expression of a functional GLP-1R in the liver of rodent models.

The ectopic GLP-1R was activated by endogenous proglucagon products in lean mice (Supplemental figure 1) where it served to increase blood glucose in a variety of experimental paradigms, functionally recapitulating GcgR agonism. In GLP-1R KO mice, the ectopic hepatic receptor increased glucose excursion in response to semaglutide (Figure 1). This effect was not seen in control GLP-1R KO mice and is the opposite response one would expect to see from semaglutide in a WT animal. Finally, we see ectopic GLP-1R agonism drive an increase in energy expenditure in response to GLP-1R and dual GLP-1R/GIPR agonists, an effect not normally associated with these compounds [8]. This is interesting insofar as simply expressing a foreign Gαs-coupled GPCR in the hepatocyte and activating that receptor with a suitable ligand appears to be sufficient to replicate much of the pharmacology of GcgR *in vivo* without the deleterious effects of GcgR pharmacotherapy on HR. It appears the hepatocyte has the necessary machinery to express the GLP-1R, transport it to the cell membrane, and enable signal transduction, as we did no modification of the *mGlp1r* gene to facilitate these functions. This speaks to the notion that this approach is generalizable to other cell signalling systems much like DREADD technology. Thus, cell specific ectopic expression of receptors may offer an novel platform to drive a specific cell signalling cascade in addition to the targeting approaches currently being developed across the pharmaceutical industry.

Second, agonism of the ectopic, hepatic GLP-1R creates a novel mechanism of drug clearance and may have effects on biodistribution. Endogenous GLP-1R does not regulate semaglutide pharmacokinetics and semaglutide clearance is increased with ectopic GLP-1R expression. It is interesting that this clearance mechanism can be overcome by introducing a GLP-1R ligand that displays low ligand-mediated receptor internalization. It is increasingly appreciated that GLP-1R agonist with a low receptor internalising profile[27-29], including tirzepatide[25], provide superior weight loss and glucose lowering efficacy. However, the ectopic GLP-1R expression system presented here offers another application for GLP-1R agonists with this particular *in vitro* profile where it helps reduce drug clearance by the receptor. Additionally, these data indicate that agonists which induce low receptor internalization may prove particularly useful in systems where endogenous receptor mediated drug clearance is in play.

Third, agonism of the ectopic, hepatic GLP-1R with either semaglutide or the dual GLP-1R/GIPR agonist does not increase HR over that seen with the agonist alone. The tachycardic effect of semaglutide is decreased in *Glp1r* AAV treated animals compared to controls, however this occurs at doses which do not induce superior weight loss and is likely the simple result of drug clearance by the ectopic GLP-1R. Yet, in the dual GLP-1R/GIPR agonist studies we see no additional increase in HR in the *Glp1r* AAV treated mice above peptide alone at doses that induce EE and improve weight loss. It has been posited that increased EE observed with glucagon pharmacology is inherently linked to increased HR[26], however these data demonstrate that EE and HR increase can be decoupled. In fact, the decreased tachycardic effect of semaglutide in *Glp1r* AAV treated mice suggests it is circulating exposure of the agonist that is more closely linked to these effects, rather than a secondary phenomenon driven by EE. One could hypothesize that the ectopic receptor approach which increases the WL effect of semaglutide while mitigating the transient tachycardia could amplify the known CV protective effects of the drug by decreasing stress on the heart itself and reducing peripheral arterial resistance via weight loss.

In conclusion these data support the development of a new methodology to safely introduce poly-pharmacology for the treatment of obesity. The ectopic, hepatic expression of GLP-1R and agonism of this receptor using semaglutide, Sema584, or a dual GLP-1R/GIPR agonist imbues these molecules with a novel EE enhancing efficacy that improves weight loss with no additional increase in HR. This approach of ectopically expressing and activating receptors with clinically validated pharmacophores represents a novel mechanism for engaging polypharmacy to treat obesity and perhaps other indications.

## Figure legends

<u>**Supplemental figure 1:** Lean WT mice do not show a baseline body-weight phenotype in response to the ectopic *Glp1r*.</u> Either lean, chow fed WT (A-D) or proglucagon KO (E-H) mice were injected with the *Glp1r* expressing AAV (10^12^ GC) on d0. (A,E) Body weight change in response to the AAV alone (B,F) I.P. glucose tolerance test, (C,G) oral glucose tolerance test, (D,H) or a mixed meal tolerance test blood glucose values. All data are presented as mean ± SEM.

<u>**Supplemental figure 2:** An LNP + mRNA construct successfully confers GLP-1R activity in hepatocytes of GLP-1R KO mice</u> (A) *In vitro* cAMP accumulation induced by semaglutide in cells transfected with GLP-1R via LNP. *In vivo* stimulation of blood glucose in GLP-1R KO mice with GLP-1R reintroduction via LNP delivered by either (B) intravenous or (C) intraperitoneal injection. All data are presented as mean ± SEM.

<u>**Supplemental figure 3:** Endogenous GLP-1R does not modify the circulating PK profile of semaglutide.</u> Circulating semaglutide profile in DIO WT and GLP-1R KO mice dosed acutely with semaglutide at 3 doses (3, 10, 30 nmol/kg); all data are presented as mean ± SEM.

