## Supplementary figures and images for "Ectopic, hepatic GLP-1R agonism enhances the weight loss efficacy of GLP-1 analogues"

### Supplemental figure 2

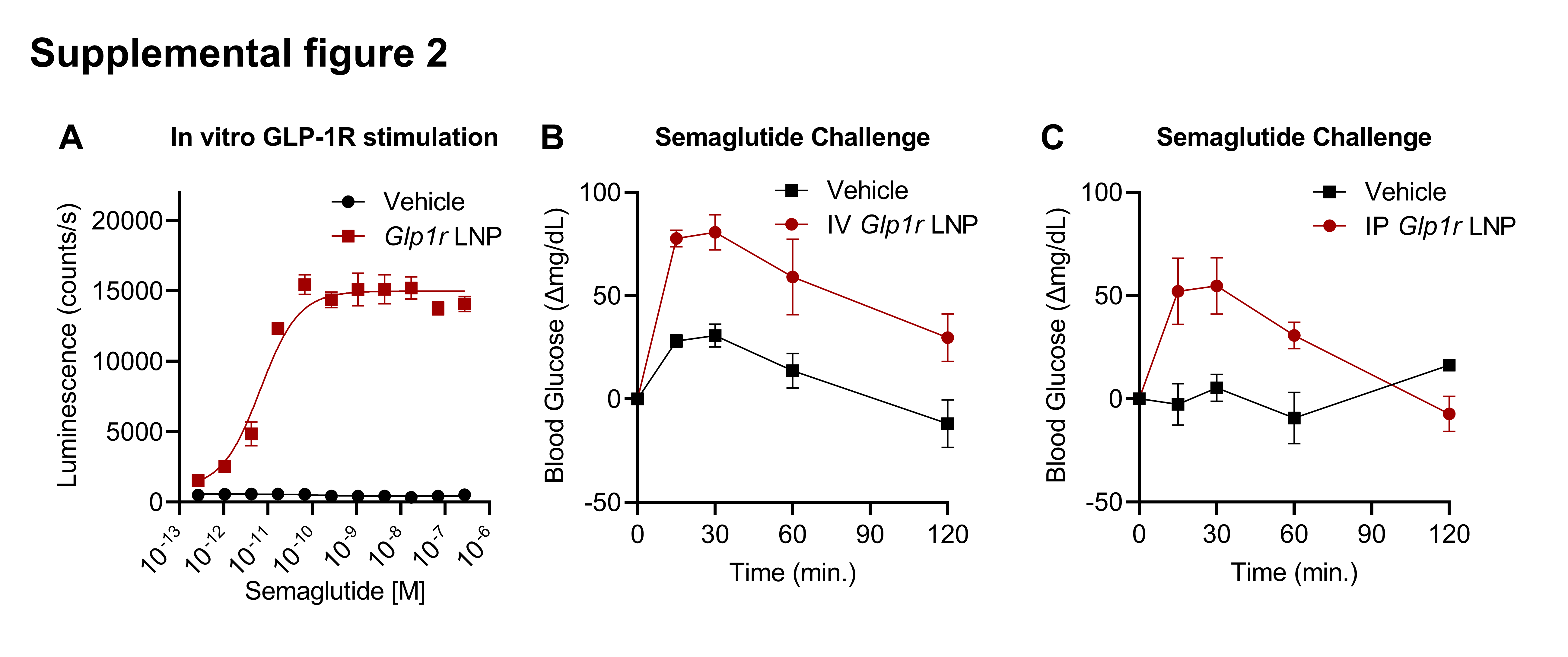

### Supplemental figure 3

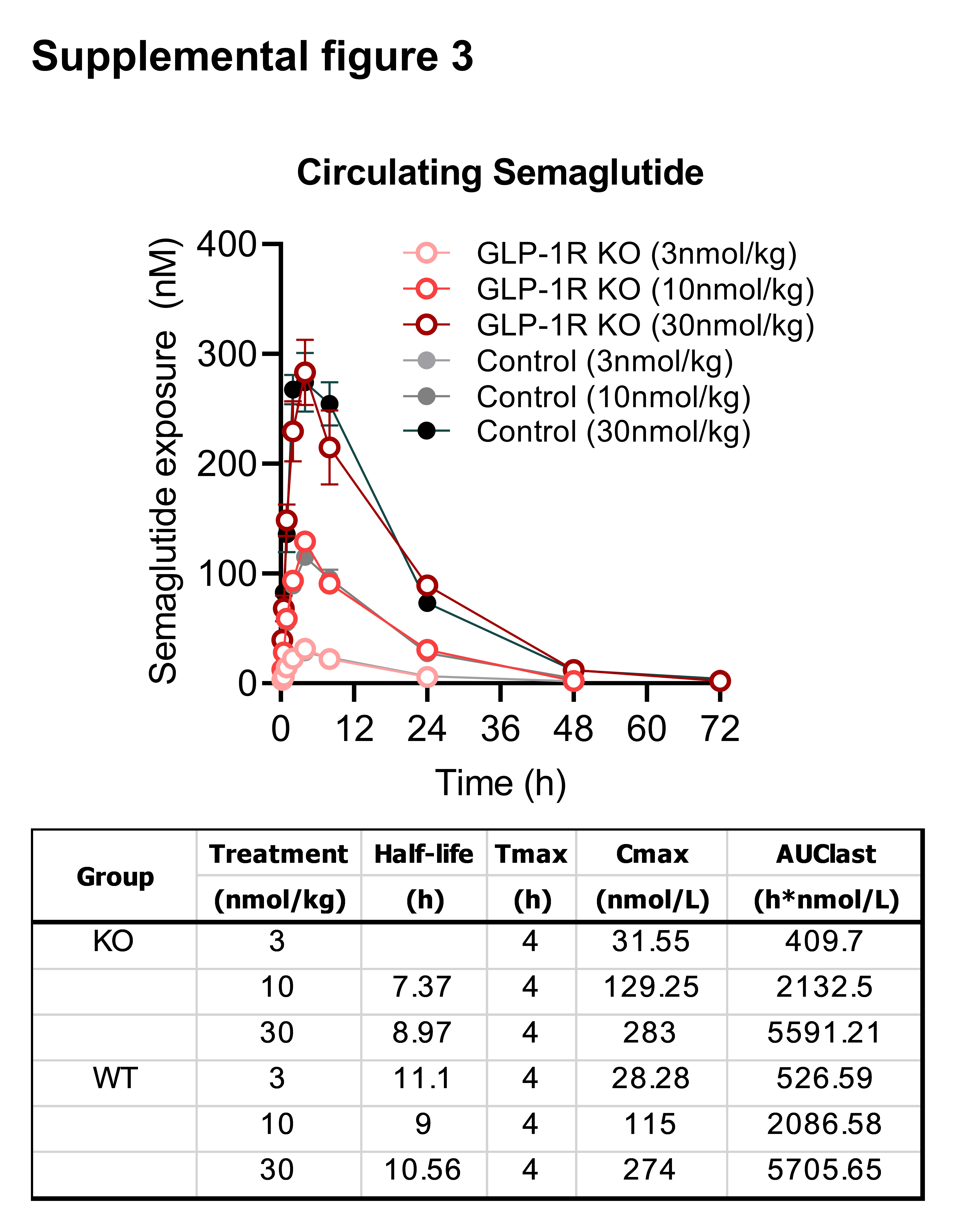

### Supplmental figure 1

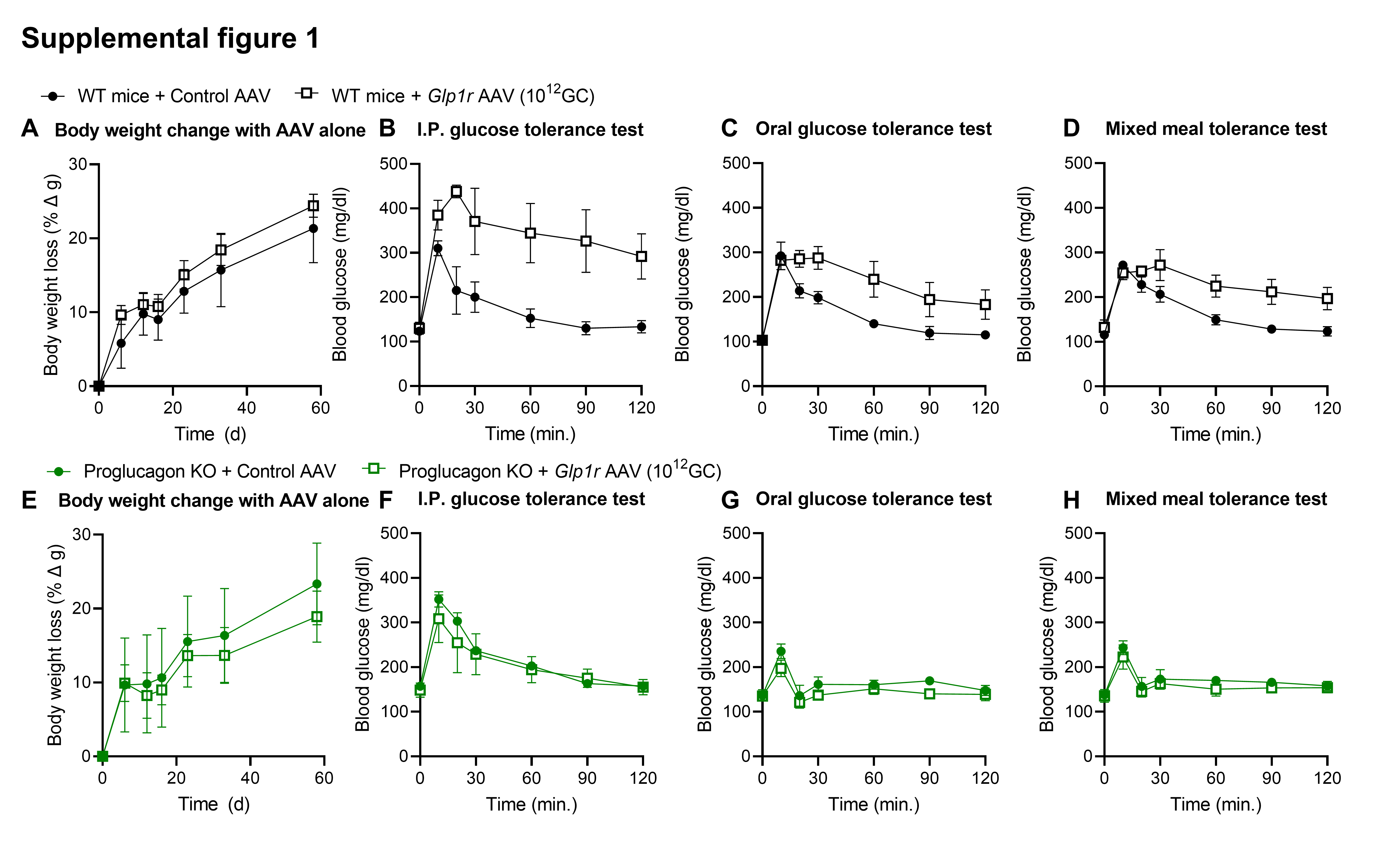
